# People Can Create Iconic Vocalizations to Communicate Various Meanings to Naïve Listeners

**DOI:** 10.1101/148841

**Authors:** Marcus Perlman, Gary Lupyan

## Abstract

The innovation of iconic gestures is essential to establishing the symbolic vocabularies of signed languages, but might iconicity also play a role in the origin of various spoken words? Can people create novel vocalizations that are comprehensible to naïve listeners without the use of prior conventions? To test this capacity, we launched a contest in which participants submitted a set of non-linguistic vocalizations for 30 meanings spanning actions, humans, animals, inanimate objects, properties, quantifiers and demonstratives. The winner – who received a monetary prize – was judged by the ability of naïve listeners to successfully infer the meanings of the vocalizations. We report the results from the contest, along with a series of experiments and analyses designed to evaluate the vocalizations for: 1) their comprehensibility to naïve listeners; 2) the degree to which they resembled their meanings, i.e., were iconic; 3) agreement between producers and listeners in what constitutes an iconic vocalization; and 4) whether iconicity helps naïve listeners learn the vocalizations as category labels. The results show that contestants were able to create iconic vocalizations for a wide array of semantic domains, and that these vocalizations were largely comprehensible to naïve listeners, as well as easier to learn as category labels. These findings provide a compelling demonstration of the extent to which iconic vocalizations can enable interlocutors to establish understanding through vocalizations in the absence of conventions. This suggests the possibility that, prior to the advent of full-blown spoken languages, people could have used iconic vocalizations to ground a spoken vocabulary with considerable semantic breadth.

## Introduction

In the parlor game charades, players are challenged to shed their spoken language and communicate using gestures. To succeed in creating a signal that successfully communicates the intended meaning to their teammates, players typically make use of *iconicity* – a resemblance between the form of a signal and its meaning. Iconicity enables a receiver to gain some understanding of a signal without relying on a previously learned conventional form-meaning association. In this way, iconicity can ground a new signal and imbue it with meaning.

Outside of the parlor, people face a similar challenge when communicating with someone who speaks a different language, a situation in which iconic gestures can likewise serve to help understanding^1^. With extended interactions, iconic gestures – along with indexical gestures like pointing – can support the formation of more fully-fledged gestural symbol systems. For example, deaf children with hearing parents use iconic gestures as the basis for more symbolic homesign systems ^2,3^. And within predominantly deaf communities that originally lack a common language, iconicity plays a crucial role in the emergence of full signed languages ^4–6^.

While the importance of iconicity to the birth of signed languages is clear, its role in spoken languages is much less so, in part because spoken languages are so ancient. In this study, we examined the possibility that the words of spoken languages may have been formed in a parallel way to the creation of many signs—through a process rooted in iconicity, but in the vocal rather than visual modality. To do this, we tested the extent to which people are capable of creating non-linguistic vocalizations that are effective at communicating various meanings to naïve listeners. We launched a contest— *The Vocal Iconicity Challenge—* in which participants were challenged to communicate a set of basic meanings by inventing novel vocalizations. We assessed the winner of the contest – motivated by a $1000 prize – by the ability of naïve listeners to successfully infer the meanings of the created vocalizations.

## Background

### Iconicity in speech and vocalization

In contrast to the clear influence of iconicity in gesture and sign, many have argued that speech affords a very limited potential for iconicity ^1,4,7,8^. For instance, Tomasello (2008) observed that it is difficult to imagine people inventing “vocalizations to refer the attention or imagination of others to the world in meaningful ways – beyond perhaps a few vocalizations tied to emotional situations and/or a few instances of vocal mimicry” ^1^. Similarly, Pinker and Jackendoff (2005: p. 209) noted that “Most humans lack the ability (found in some birds) to convincingly reproduce environmental sounds.” They proposed that the human capacity for vocal imitation is essentially limited to “a capacity to learn to produce speech.” Within actual spoken languages, Saussure’s ^9^ notion of the arbitrariness of the sign is commonly adopted as essential to the nature of spoken words ^10^, and the number of iconic and imitative words has been assessed as “vanishingly small” ^11^, exceptional “asterisks” to the principle of arbitrariness^12^. According to Hockett ^8^, this is the inevitable consequence of the limited dimensions of speech to afford iconicity.

Some scholars of language evolution have pointed to claims like these – postulating limited potential for iconicity in vocalization and speech – as support for the theory that the first languages were gestural. On this idea, the first languages must have originated from iconic gestures that served, eventually, to scaffold arbitrary vocalizations ^1,3,4,13,14^. Similar rationale supports an argument for a multimodal division of labor in which gestures and speech co-evolved, but with gesture carrying the iconic load and bootstrapping arbitrary speech ^16^.

However, an improved understanding of the vocabularies of non-European languages and increased empirical scrutiny of the “arbitrariness of the sign” dogma have revealed that iconicity in spoken languages is much more pronounced than previously suspected ^17–20^. For example, many spoken languages have substantial inventories of ideophones (also called mimetics^21^, expressives^22^, or phonaesthemes^23^), a distinctly iconic class of words used to express sensory meanings across diverse domains like animate and inanimate sounds, manner of motion, size, visual patterns, textures, inner feelings and cognitive states^24,25^. Some languages, such as Japanese, can contain thousands of these depictive words^18^. Linguists have also identified many iconic words outside of the ideophone lexical class. For example, across many languages, words expressing *smallness* and related concepts feature high front vowels, while *large* concepts feature low back vowels ^26,27^. This pattern may help explain the differences between typically feminine and masculine personal names ^28^. It may also motivate the forms of the indexical words used to refer to proximal and distal referents, such as the translational equivalents of the English demonstratives “here” and “there” ^29^ and “this” and “that” ^29,30^. Words for proximal referents tend to contain front vowels, whereas distal words tend to contain back vowels. Iconicity is also prevalent in some anatomical vocabulary, as languages show a prevalence of nasal consonants in words for “nose” and bilabial consonants for “lip” ^31^. Recent large-scale analyses of basic vocabulary across thousands of languages have confirmed the prevalence of some of these iconic relationships between forms and meanings ^32^.

In addition to iconicity in the phonology of words, a more dynamic form of iconicity in spoken language is found in the intonation, tempo, and loudness – the prosody – of speech. Bolinger ^33^ suggested that a fundamental function of prosody, especially intonation, is the iconic expression of emotion. Ohala ^34^ and others ^35^ have noted that speakers also use prosody to express qualities related to size, dominance, and strength. More broadly, experimental evidence indicates that prosody can enhance the iconicity of ideophones spanning meanings across the senses ^36^.

Speech production experiments have also shown that speakers sometimes produce iconic modulations in their prosody when communicating about a range of meanings. For example, speakers have been shown to increase or decrease their tempo when respectively describing a fast or slow-moving event, and to raise or lower their pitch when describing upward or downward movement or when referring to something small or large ^37,38^. Iconic prosody may be especially evident in speech directed towards young children. Three adults were asked to produce novel words in infant directed speech, with the meaning paired to one of 12 antonymic adjectives ^39^. Analysis of the utterances showed certain consistent differences between the prosodic properties associated with particular meanings, including properties like fundamental frequency, amplitude, and duration. For instance, *strong* was expressed with higher amplitude than *weak*; *happy* with higher pitch, higher amplitude, and a shorter duration than *sad*; and *tall* with a longer duration than *short*. Moreover, naïve listeners were better than chance at selecting a picture matching the original meaning of the word from two alternatives ^38^.

### Inventing novel vocalizations

While the human ability to vocally imitate is often assumed to be poor ^10^, empirical results paint a different picture. Lemaitre and Rocchesso ^40^ asked participants to imitate various mechanical and synthesized sounds or to provide verbal descriptions of them. When these were played back to listeners, participants were better at identifying many of the original sounds from the vocal imitation than from the verbal description. A subsequent study found that people are effective at communicating with vocal imitations because they focus on a few salient features of the source, rather than producing a high fidelity representation ^41^.

In addition to the direct imitation of sounds, recent experiments have shown that people are able to spontaneously invent iconic vocalizations to represent various other kinds of meanings ^19^. Participants played a charades-type game in which they took turns improvising non-linguistic vocalizations to communicate meanings from 30 different antonym pairs, including contrasting words like *alive* and *dead, dull* and *sharp, hard* and *soft, fast* and *slow, bad* and *good,* and *bright* and *dark*. Their vocalizations were highly consistent in the particular acoustic features that were used to distinguish contrasting words in more than two thirds of the antonymic pairs. For example, *rough* compared to *smooth* was expressed with aperiodic sounds marked by a lower harmonics-to-noise ratio, *small* compared to *large* with quiet, high-pitched sounds, and *fast* compared to *slow* with loud, high-pitched, quickly repeated sounds.

Other studies have shown that these invented vocalizations are, to some degree, understandable to naïve listeners. One experiment compared the use of non-linguistic vocalization and gesture to communicate 18 items that included emotions (e.g. *disgust*, *tired*), actions (e.g. *throwing*, *chasing*) and objects (e.g. *predator*, *tree*) ^42,43^. Compared to the chance rate (5.6%), accuracy in the initial block was highest for emotions (~60%), next for actions (~40%), and lowest for objects (~10%). In another study, participants took turns for ten rounds producing non-linguistic vocalizations to communicate a set of meanings from nine antonymic pairs of words, including items like *bad, good, big*, *small, down*, *up, far*, *near, fast*, *slow, few*, *many, rough*, and *smooth* ^44^. With few exceptions, each meaning was expressed with characteristic acoustic properties that distinguished it from each other meaning. In subsequent playback experiments, naïve listeners were better than chance at guessing all but one of the 18 meanings, and for 15 of them, their accuracy was at least 20% and as high 73%, compared to a chance rate of 10%.

## Current Study

The work reviewed above shows that (1) iconicity pervades spoken languages much more than previously realized and (2) that people have some ability to invent iconic vocalizations that can be understood by naïve listeners. But just how good are people at doing this? What is the full extent to which humans can ground a symbol system through iconic vocalizations, without the use of gesture? We investigated this upper limit by conducting the *Vocal Iconicity Challenge!* – a contest that challenged participants to create iconic vocalizations for 30 meanings that spanned an array of semantic domains, including actions, humans, animals, inanimate objects, properties, quantifiers and demonstratives. The iconicity of the vocalizations was evaluated by the ability of naïve listeners to guess their meanings from a set of alternatives. To push the level of motivation, the winning team or individual whose vocalizations were guessed most accurately received a prize of $1000. Submissions included participants affiliated with several prominent linguistics and language evolution programs at universities in the United States and Europe.

## Results

Here we report the results from the contest, along with a series of experiments and analyses designed to evaluate the vocalizations for 1) their comprehensibility to naïve listeners; 2) the degree to which they resembled their meanings, i.e., were iconic; 3) agreement between producers and listeners in what constitutes an iconic vocalization; and 4) whether iconicity helps naïve listeners learn the vocalizations as category labels. The overarching goal of the analyses was to assess the capacity for people to use iconic vocalizations to ground a spoken symbol system.

First, we analyzed the comprehensibility of the vocalizations by asking naïve listeners to guess their meaning from a set of alternatives. The vocalizations were presented to listeners in two testing conditions (see Methods). In within-category testing, listeners selected the meanings from alternatives in the same broad semantic category (actions, properties, nouns), and in between-category testing, the meanings were selected from alternatives from across the three main semantic categories, including some potentially confusable alternatives (e.g. *rock, pound, dull*). For each submitted set of vocalizations, we calculated the guessing accuracy in the two conditions. Additionally, we examined guessing accuracy for the different meanings and semantic categories, as well as the errors that guessers made.

Comprehensibility provides one index of iconicity, but we also wanted to assess the degree to which listeners actually perceived a resemblance between form and meaning. Therefore, we next asked naïve listeners to directly rate the degree to which the vocalizations sounded like their intended meaning. We determined whether vocalizations for some meanings tended to be rated as more iconic than others, and also how well the iconicity ratings predicted the ability of listeners to correctly guess their meanings.

Third, we wanted to examine iconicity from the perspective of creating and articulating the vocalizations, particularly whether the level of agreement between producers correlated with the guessing accuracy of listeners. We did this by measuring the consistency between contestants in the vocal qualities they used – fundamental frequency, duration, voice quality, and loudness – to represent each meaning. We then tested whether the level of agreement in how to produce an iconic vocalization for a given meaning correlated with the guessing accuracy of naïve listeners.

Finally, we examined whether iconicity helps people learn the vocalizations as labels for categories. Based on the iconicity ratings, we used low-, medium-, and high-iconicity vocalizations as stimuli to test whether naïve listeners were better at learning the meanings of more iconic signals. We manipulated the feedback that learners received – full feedback indicating the correct response, or accuracy only – to assess whether iconicity might be especially helpful under more challenging learning conditions.

### Comprehensibility of Vocalizations

Figure 1 shows box and whisker plots of accuracy in within- and between-category testing for each of the 11 submissions. To test whether accuracy over all submissions was higher than chance, we constructed logistic mixed effects models of accuracy, with the intercept offset to the chance level, and random intercepts for listener, meaning, and submission ID. Similarly, we constructed models of accuracy to compare specific submissions to chance (except without submission ID as a random intercept). In within-category testing, average accuracy over all submissions was 39.0% (bootstrap 95% CI = [36.2%, 41.9%]), significantly higher than chance (10%), *b*_*0*_ = 1.64, 95% CI *=* [1.11, 2.16], *z* = 6.47, *p* < 0.0001. Accuracy ranged from 58.0% for the top submission to 20.3% for the last-place submission, which was still reliably higher than chance, *b*_*0*_ = 0.62, 95% CI = [0.057, 1.04], *z* = 2.55, *p* = 0.011. In between-category testing, average accuracy was 35.9% (bootstrap 95% CI = [0.35, 0.40]), significantly higher than chance (10%), *b*_*0*_ = 1.48, 95% CI = [0.94, 2.02], *z* = 5.69, *p* < 0.0001. Accuracy ranged from 56.0% for the winning submission to 12.2% for the last-place submission. The last-place submission was not higher than chance, but the next-to-last place submission (21.5%) did reliably exceed chance, *b*_*0*_ = 0.71, 95% CI = [0.16, 1.14], *z* = 2.99, *p* = 0.0028. To test whether there was a difference in accuracy between within-category and between-category testing, we constructed a logistic mixed effects model of accuracy, with testing condition as a fixed effect. Random intercepts were included for meaning, submission ID, and listener, and random slopes of category were included for meaning and submission ID. The model failed to indicate a significant difference in accuracy between the testing conditions, *b* = 0.17, *z* = 1.56.

**Figure 1.**
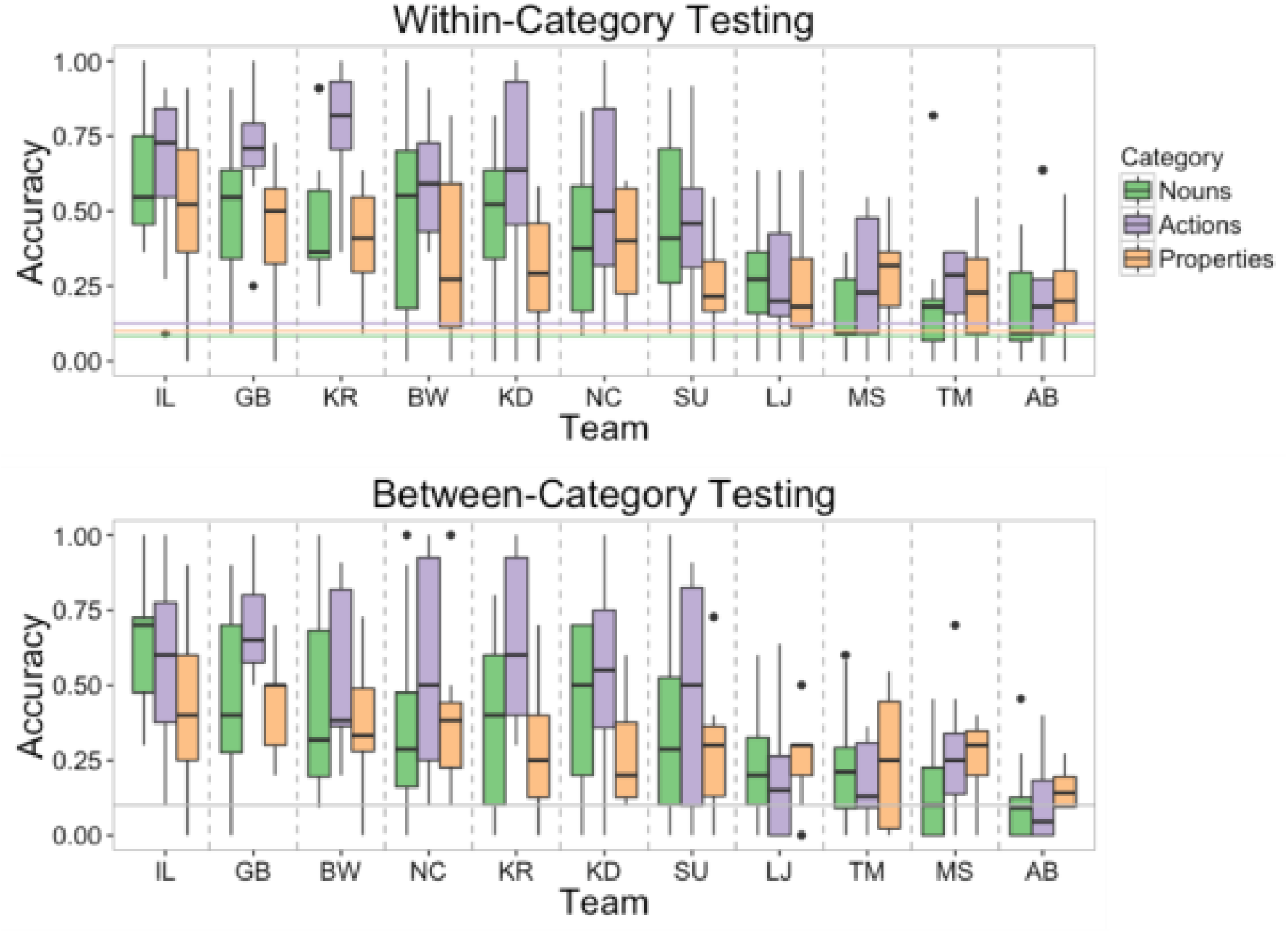
Box and whiskers plots of accuracy for each submission. The line within each box signifies the median, the bottom and top hinges of the box correspond to the first and third quartiles, the bottom and top lines extending from the box correspond to 1.5 * IQR (inter-quartile range), and the points to outliers beyond this range. Top plot shows accuracy in within-category testing. The two dashed horizontal lines indicate chance accuracy, with color matched to testing round. In within-category testing, chance was 8.3% for the 12 nouns, 12.5% for the 8 actions, and 10% for the 10 properties. Chance for between-category testing was 10% for the 10 alternatives.

Figure 2 shows box and whisker plots of accuracy for each meaning under both testing conditions. Averaged across the two conditions, accuracy for individual items ranged from 15.4% (bootstrap 95% CI = [11.5%, 20.0%]) for “that” to 72.7% (bootstrap 95% CI = [57.3%, 87.0%]) for “sleep”. For each of the 30 meanings, we constructed logistic regression models of accuracy, with the intercept offset to the chance level, and with participant and submission ID as random intercepts. These models showed that of the 30 meanings, 27 were guessed more accurately than chance, *b*_*0*_’s > 0.92, *z*’s > 2.72, *p*’s < 0.007. “Fruit” was guessed correctly at a rate marginally above chance, *b*_*0*_ = 0.65, 95% CI = [-0.02, 1.33], *z* = 1.90, *p* = 0.057. Only “that” and “gather” were not guessed significantly above chance, *b*_*0*_’s < 0.34, *z*’s > 0.65.

**Figure 2.**
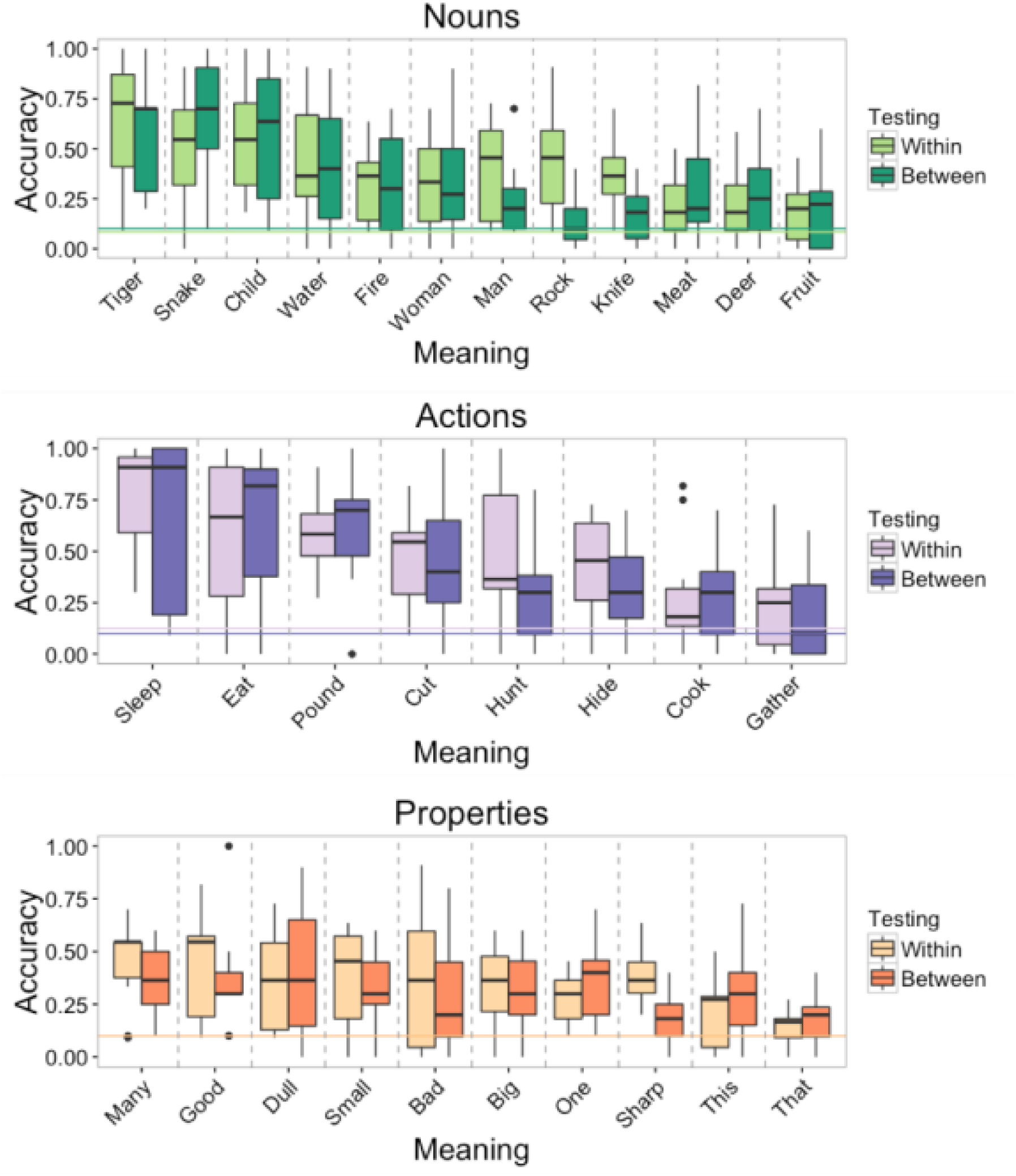
Box and whisker plots of accuracies for each meaning by category. The line within each box signifies the median, the bottom and top hinges of the box correspond to the first and third quartiles, the bottom and top lines extending from the box correspond to 1.5 * IQR (inter-quartile range), and the points to outliers beyond this range. Top plot shows nouns, the middle plot actions, and the bottom plot properties. The two dashed horizontal lines indicate chance accuracy, with color matched to testing round. In within-category testing, chance was 8.3% for the 12 nouns, 12.5% for the 8 actions, and 10% for the 10 properties. Chance for between-category testing was 10% for the 10 alternatives.

Across both testing phases, the accuracy for actions was 45.6% (bootstrap 95% CI = [41.0%, 50.1%]), for properties 31.8% (bootstrap confidence interval = [28.6%, 34.3%]), and for nouns 36.6%, (bootstrap 95% CI = [33.0%, 40.6%]). For each condition, we constructed a logistic mixed effects model to assess the relationship between guessing accuracy and the meaning categories. The model included random intercepts for listener, submission ID, and meaning. For the within-category condition, category was included as a random slope for submission ID, and for the between-category condition, category was included as random slopes for listener and submission ID. In within-category testing – in which vocalizations for actions were selected from the 8 alternatives in the set of actions, properties from the 10 properties, and nouns from the 12 nouns – actions were guessed with marginally higher accuracy than properties, *b* = −0.74, 95% CI = [-1.23, 0.13], *z* = - 1.90, *p* = 0.058, but not higher than nouns, *b* = −0.55, *z* = −1.58. In between-category testing, in which all items were selected from 10 alternatives, the meanings of actions were not guessed more accurately than properties, *b* = −0.50, *z* = −1.19, or nouns, *b* = - 0.37, *z* = −0.95.

We next examined whether vocalizations for particular meanings were more likely to be confused with some meanings rather than others. Figure 3 shows confusion matrices for guessing in the two testing conditions. The items are ordered according to semantic similarity based on Google’s *word2vec* semantic vectors (see Methods). The warm-colored diagonals from upper left to bottom right show that listeners most frequently selected the intended meaning of the vocalizations. However, the matrices reveal that some meanings were often confused. For instance, in within-category testing (Figure 3 A-C), vocalizations for “woman” were often confused with “child”, “that” was confused with “dull”, and “many” with “bad”. The between-category matrices (Figure 3 D-F) show a tendency for participants to confuse semantically related meanings between categories, such as “knife” with “cut” and “child” with “small”.

**Figure 3.**
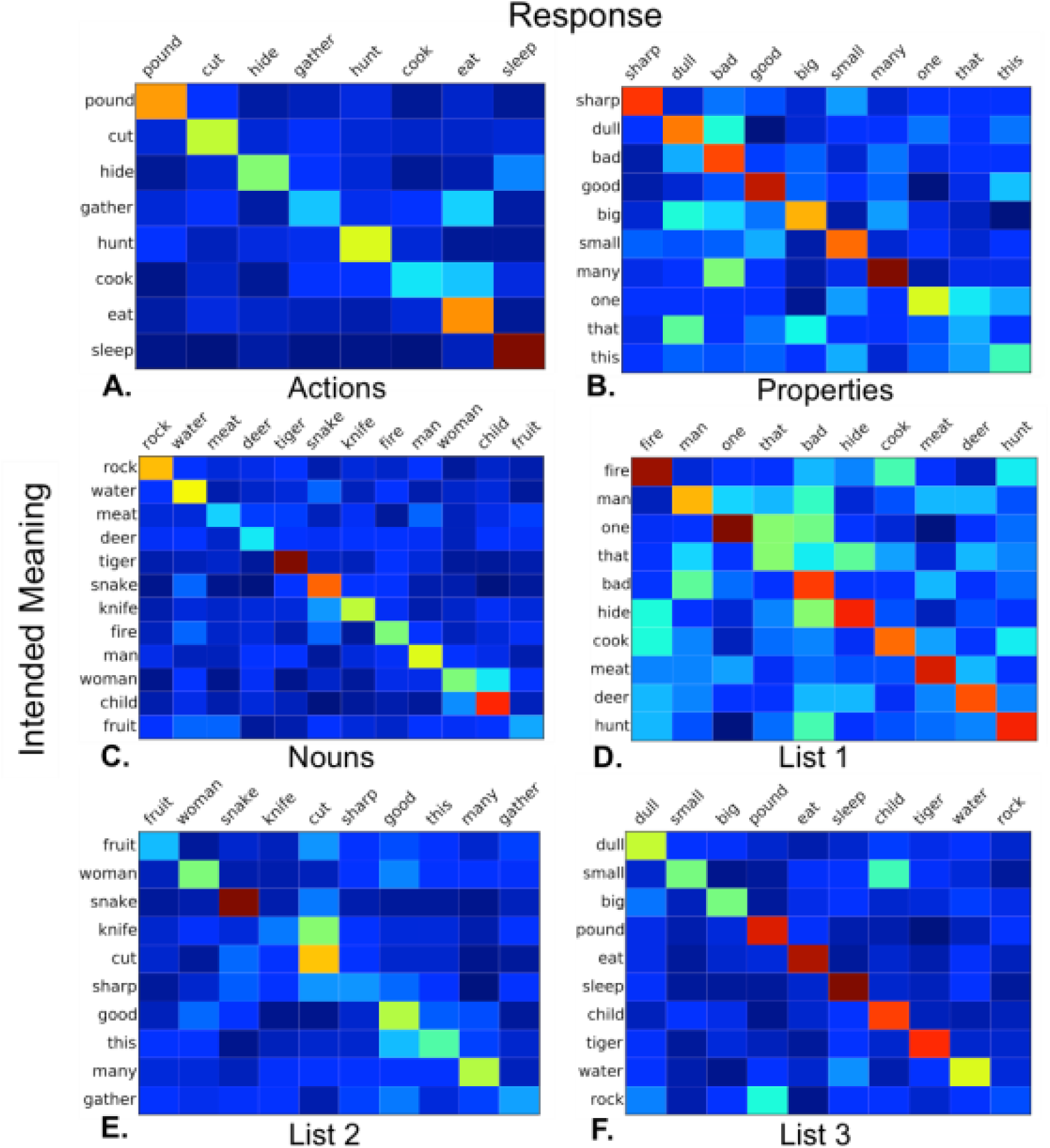
Confusion matrices showing results from the two testing conditions. The y-axis shows the intended meaning of the vocalizations, and the x-axis shows the guessed meanings. The warmer the color of a cell, the more frequently that response was confused for that intended meaning. Items are ordered according to semantic similarity based on the correlation between their *Word2Vec* semantic vectors (see Methods). *A* shows actions, *B* Properties, *C* nouns, *D* list 1, *E* list 2, and *F* list 3. Intended meanings are on the y-axis, and responses are on the x-axis.

We then tested whether the semantic similarity between the intended meaning of the vocalization and the selected meaning (based on Google’s *Word2Vec* semantic vectors) correlated with the proportion of trials in which listeners confused these meanings. A Pearson’s correlation test indicated a significant positive relationship between these variables, *r* = 0.14, 95% CI = [0.05, 0.23], *t*(463) = 3.00, *p* = 0.003. (When correct responses were included with a similarity index of 1, the correlation increases to *r* = 0.71, *t*(493) = 22.4, 95% CI = [0.66, 0.75], *p* < 0.001). Next we constructed a linear mixed effects model of the proportion of confusions between two meanings with their semantic similarity as a predictor. Random intercepts were included for meaning and response, and random slopes of similarity were included for meaning and response. With correct responses excluded, the model showed that semantic similarity was a significant predictor of the proportion of confusions, *b* = 0.065, 95% CI = [0.20, 0.11], χ1^2^ = 4.47, *p*= 0.0063. (When correct responses were included, semantic similarity was a much stronger predictor, *b* = 0.29, 95% CI = [0.24, 0.34], χ1^2^ = 58.4, *p* < 0.001.)

### Ratings of form-meaning resemblance

To measure the iconicity of the vocalizations, we asked naïve listeners to rate the degree to which they “sound like” their intended meaning, thereby providing a more direct evaluation of form-meaning resemblance (cf. ^45^, which used this method to evaluate the iconicity of English and Spanish words). Figure 4 shows the distribution of iconicity ratings for each meaning by semantic category. To determine whether the level of iconicity of the vocalizations differed between semantic categories, we constructed a mixed effects model of iconicity rating, with semantic category as a fixed effect. Vocalization, submission ID, and listener were modeled as random intercepts. Random slopes of category were included for submission ID and listener. Model comparisons showed a marginally reliable effect of semantic category on ratings, χ1^2^ = 4.73, *p* = 0.094. A subsequent model comparing just actions and nouns showed that actions were rated significantly higher in iconicity, *b* = −0.46, 95% CI = [-0.90, −0.02], χ1^2^ = 4.07, *p* = 0.044.

**Figure 4.**
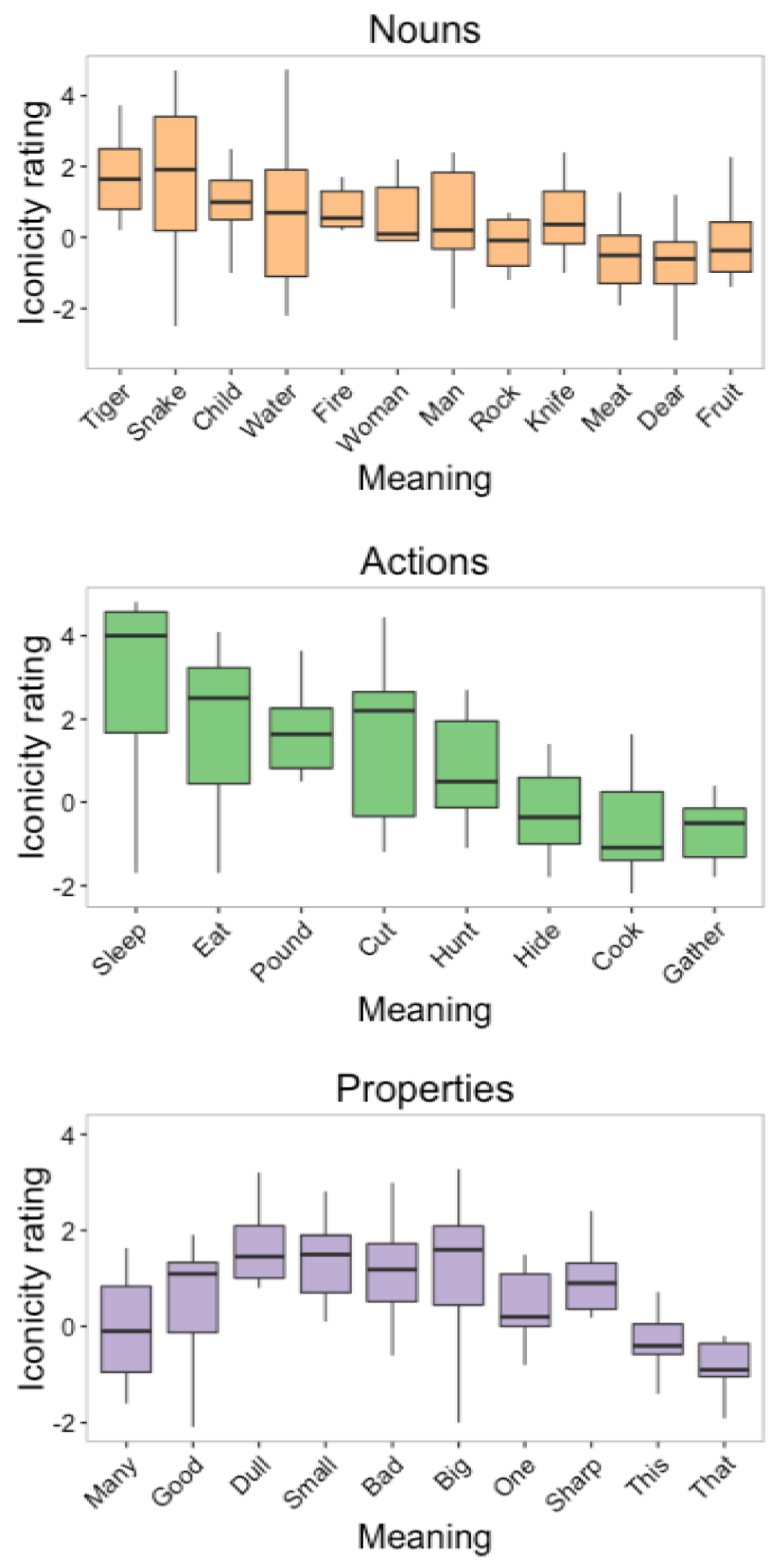
Box and whisker plots of iconicity ratings for each meaning. The line within each box signifies the median, the bottom and top hinges of the box correspond to the first and third quartiles, the bottom and top lines extending from the box correspond to 1.5 * IQR (inter-quartile range), and the points to outliers beyond this range. Meanings are ordered from lowest to highest guessing accuracy. This ordering shows that iconicity ratings and guessing accuracy were highly correlated, especially for nouns and actions.

We then used a logistic mixed effects model to determine whether the iconicity ratings were a reliable predictor of guessing accuracy. This model included the mean iconicity rating for a vocalization as a main effect. Random intercepts were modeled for listener, submission ID, and vocalization. Random slopes of iconicity rating were included for listener and submission ID. Figure 5A shows the fit of this model against a scatter plot of accuracy as a function of iconicity rating. Iconicity rating was a reliable predictor of guessing accuracy, *b* = 0.54, 95% CI = [0.44, 0.63], *z* = 10.86, *p* < 0.001; R^2^ = 0.15 for fixed effects and R^2^ = 0.35 for fixed and random effects.

**Figure 5.**
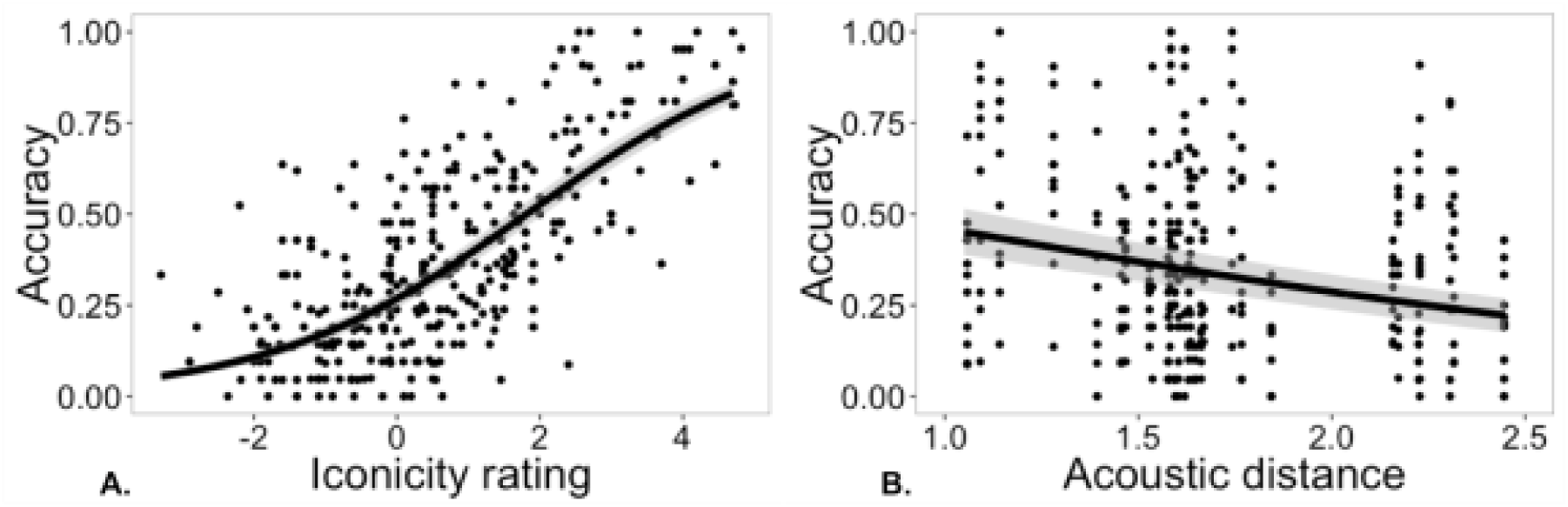
A. Scatter plot of guessing accuracy for each vocalization as a function of its iconicity rating. The line is the fit of a logistic mixed effects model and +/-1SE of the model estimate. B. Guessing accuracy as a function of the acoustic distance of each meaning. Each point represents a vocalization, but the acoustic distance is for each meaning, resulting in the vertical series of points. The line shows the fit of a logistic mixed effects model, and the grey band indicates the standard error of the model.

### Agreement between producers and listeners

We also examined the iconicity of the vocalizations from a production standpoint. For each meaning, we computed the acoustic distance between the vocalizations from each submission based on fundamental frequency, duration, intensity, and harmonics-to-noise ratio (see Methods). We then tested whether the degree of similarity of vocalizations (i.e. smaller acoustic distance) for a given meaning predicted the guessing accuracy of naïve listeners. We reasoned that if producers invented similar-sounding vocalizations for a meaning, then this would indicate an especially strong iconic association between form and meaning, which ought to be reflected in more accurate guessing. To test this, we constructed a logistic mixed effects model, with acoustic distance as a main effect, random intercepts for listener, submission ID, and vocalization, and random slopes of acoustic distance for listener and submission ID. Figure 4B shows the fit of this model against a scatter plot of guessing accuracy as a function of acoustic distance for each meaning. The analysis showed that acoustic distance was a reliable predictor of guessing accuracy, *b* = −0.78, 95% CI = [-1.16, −0.41], *z* = −4.10, *p* < 0.001.

When iconicity rating was added to this model as a main effect, both rating, *b* = 0.54, 95% CI = [0.46, 0.62], *z* = 13.93, *p* < 0.001, and distance, *b* = −0.38, 95% CI = [-0.68, - 0.09], *z* = −2.54, *p* = 0.011, were reliable predictors of guessing accuracy. Thus, guessing accuracy was related to both listeners’ judgments of form-meaning resemblance – a “sounds like” relationship, and to the level of agreement between the producers of the vocalizations – the degree to which they used similar vocal qualities to express each particular meaning.

### Learnability as category labels

Lastly, we conducted a learning experiment to examine whether the iconicity of vocalizations plays a role in the ability of people to learn them as category labels.

Participants (University of Wisconsin undergraduates) were tasked with learning to associate twelve vocalizations with 12 noun categories (e.g., fire, man, etc.). They were randomly assigned into a high, medium, or low-iconic group. All three groups completed the same learning task, but learned to associate the categories with vocalizations that were – according to the iconicity ratings we collected – high, medium or low in iconicity. In addition, participants were assigned to one of two feedback conditions: full feedback in which the correct response was indicated, and accuracy only feedback in which their response was only indicated as correct or incorrect.

The results of the learning experiment are shown in Figure 6. To evaluate the results, we constructed a logistic mixed effects model of guessing accuracy. The model included iconicity rating, block, and feedback as main effects, as well as terms for interactions between these variables. Random intercepts were included for subject and vocalization, as well as random slopes of block for subject and vocalization. Not surprisingly, accuracy increased over blocks, showing that participants were able to learn the categorical meanings of the vocalizations, *b* = 0.58, 95% CI = [0.51, 0.66], *z* = 15.31, *p* < 001. There was a reliable effect of iconicity on accuracy, *b* = 3.51, 95% CI = [2.45, 4.57], *z* = 6.52, *p* < 0.001, such that accuracy was highest in the high iconicity conditions (86.5%), followed by the medium iconicity conditions (58.5%), followed by the low iconicity conditions (47.3%). The model also showed that accuracy was higher in the full feedback conditions (77.4%) compared to accuracy only (50.8%), *b* = 2.42, 95% CI = [1.90, 2.94], *z* = 9.17, *p* < 0.001.

**Figure 6.**
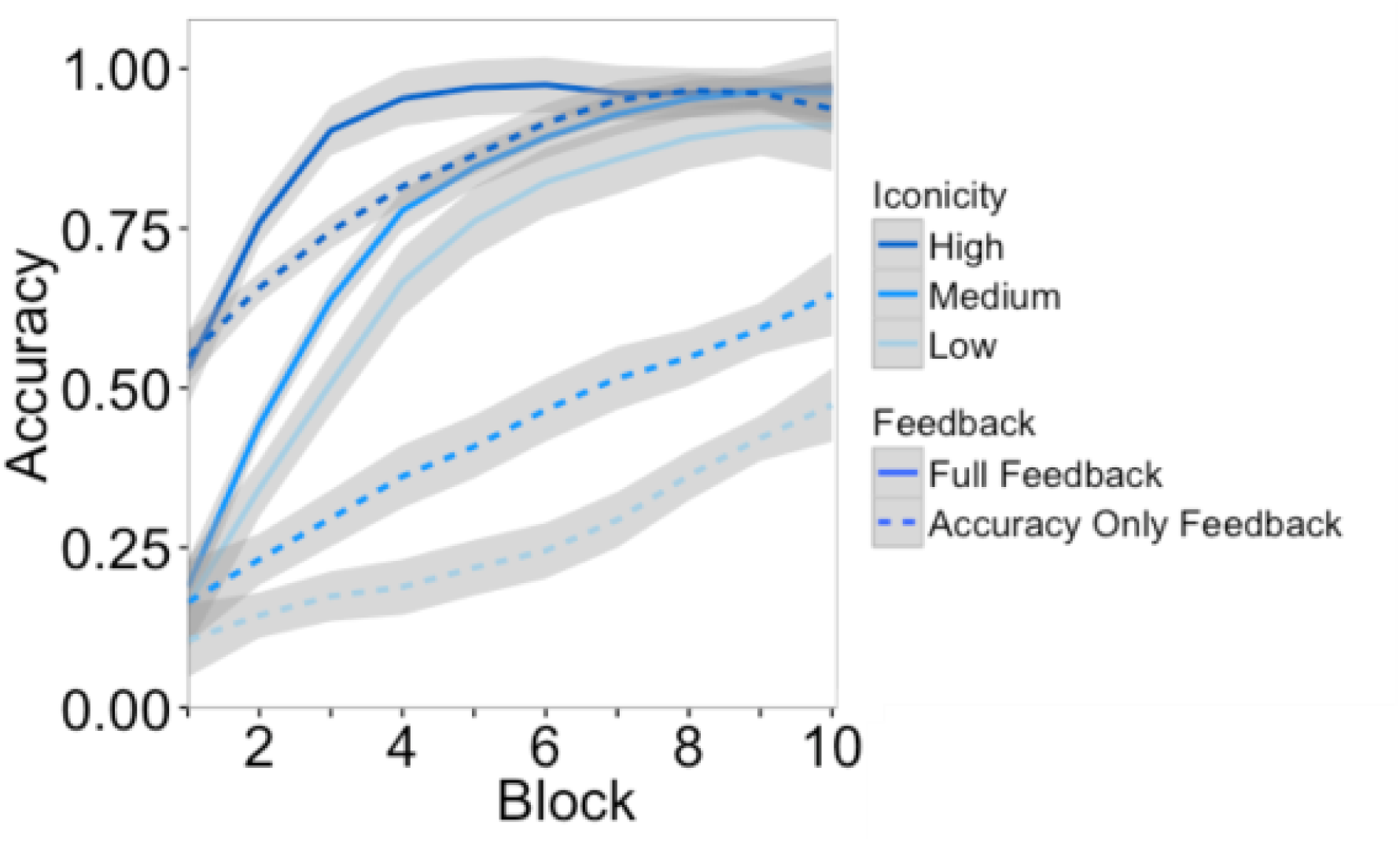
Accuracy over blocks in the learning experiment. Responses were selected from 12 alternatives, with chance-level accuracy at 0.083.

Learning was faster with full feedback than accuracy-only feedback, as supported by a reliable interaction between feedback and block, *b* = 0.40, 95% CI = [,], *z* = 6.43, *p* < 0.001. Learning was faster with higher levels of iconicity, as shown by a significant interaction between iconicity and block, *b* = 0.26, 95% CI = [,], *z* = 2.82, *p* = 0.005. There was also an interaction between iconicity and feedback, *b* = −1.61, 95% CI = [,], *z* = −2.48, *p* = 0.013, suggesting that iconicity provides a larger advantage in learning with accuracy-only feedback by helping the listener to home in on the correct meaning more quickly. The iconicity boost may have been limited in the full-feedback condition as participants reached ceiling performance after only four blocks.

## Discussion

Many theories of language evolution have assumed that people have very limited means to express meanings using novel (i.e., not already conventionalized) vocalizations. As a result, it is commonly posited that manual gestures must have played an essential role in bootstrapping the formation of spoken symbols ^1,3,4,13,14^. However, evidence from spoken languages ^17,20^, as well as from experiments ^42,44^, suggests that people are surprisingly adept in inventing and interpreting novel vocalizations via iconicity – resemblance between form and meaning. Our goal in this study was to assess the upper limit of this ability. What is the potential for humans to ground a symbol system through iconic vocalizations without the benefit of gestures?

We conducted a contest in which participants – incentivized by a monetary prize – competed to devise a set of non-linguistic vocalizations to communicate 30 different meanings to naïve listeners. Extending beyond previous studies, the meanings spanned a diverse range of semantic domains, including action verbs, nouns for people, animate and inanimate things, adjectives, quantifiers, and demonstratives. After receiving the submissions, we performed a series of experiments and analyses to evaluate the vocalizations for 1) their comprehensibility to naïve listeners; 2) the degree to which the vocalizations resembled their meanings, i.e., were iconic; 3) agreement between producers and listeners in what constituted an iconic vocalization; and 4) whether iconicity helped listeners learn the vocalizations as labels for categories. The findings showed that contestants were able to create vocalizations to successfully communicate about a wide variety of meanings. For more than half of the submissions, the guessing rate across the 30 vocalizations was greater than 40%, compared to a chance rate of 10%. Over two phases of testing, guessing accuracy was reliably higher than chance for 27 of the 30 meanings (and marginally higher for one more). Analysis of the errors guessers made showed that they tended to confuse semantically related words, indicating that a sense of the meaning was often conveyed by the vocalization even when it did not lead to the correct response.

To further analyze the iconicity of the vocalizations, we asked new listeners to rate them for the degree to which they “sound like” their meaning. These iconicity ratings proved to be a highly significant predictor of guessing accuracy for the vocalizations, suggesting that listeners relied heavily on form-meaning resemblance in making their selections. In addition, we found that when contestants produced more similar sounding vocalizations for a particular meaning, the guessing accuracy for that meaning tended to be higher. Thus, when producers largely agree on the quality of vocalization to produce for a given meaning, listeners tend to agree in interpreting that meaning from the vocalization.

Finally, we conducted a learning experiment to examine whether the iconicity of vocalizations plays a significant role in how well they can be learned as category labels. Are the meanings of more iconic vocalizations easier to learn, especially in conditions when informative feedback is limited? The results showed that vocalizations that were higher in iconicity were learned faster than those that were lower in iconicity, particularly when the feedback provided just the accuracy of the response and did not indicate the correct answer. With more iconic vocalizations, listeners were quick to learn their meaning from trial and error after only a few blocks, whereas they often failed to discover the correct meaning of less iconic vocalizations.

Overall guessing accuracy was remarkably high for many of the submissions, and the results suggest that some meanings afford a high level of iconicity (e.g. *tiger*, *eat*, *many*). However, it is also clear that some meanings afford substantially less. One interesting case is the notably low guessing accuracy for the demonstratives *this* and *that,* which fits with previous results for the spatial adjectives *near* and *far* ^44^. While these spatial meanings may not translate to iconic, easily interpretable vocalizations (but see ^30^), they are well suited to pointing gestures. This highlights the potential for iconicity to play complementary roles across modalities: some meanings may better afford iconicity in vocalization and others in gesture.

The most successful submissions – particularly the top six with accuracy rates over 40% – were affiliated with academic programs conducting research in linguistics, psycholinguistics, and language evolution. This raises the possibility that the trained intuitions of language scholars might have been useful for deriving iconic vocalizations that are most understandable to naïve listeners. Another, possibly more important factor, however, is teamwork: four of the top five submissions were created by teams (M = 4.75 participants per team). Both facts point to the possible role of deliberation and interaction in devising successful iconic vocalizations. This is consistent with previous findings with an iterated version of the vocal charades task, which found that participants produced vocalizations that were more understandable to naïve listeners after repeated interactions^44^. Thus, the full expressive potential of vocal iconicity may not be spontaneously available to communicators, but may be sharpened by deliberation and interaction.

One point of qualification of our findings is that the study was limited to English speakers – both in the contestants who produced the vocalizations and in the participants who listened to them. Yet, compared to many psycholinguistic experiments, our contestants were fairly heterogeneous – from across the US, as well as from the UK and Poland, including two submissions from native German speakers and one from native Polish speakers. Is it possible that listeners relied not on iconicity, but on arbitrary conventions shared among the English-speaking population? To a degree, this concern is mitigated by the contest rules, which did not allow spoken emblems or onomatopoeia. Additionally, the findings that the iconicity ratings were a strong predictor of guessing accuracy and learning suggest that people were tuned in to the resemblance between form and meaning, and that this resemblance played a role in their performance. Nevertheless, future research is required to examine any cultural variability in the patterns we observed here.

Combined, the findings from our contest provide one of the most compelling demonstrations to date of how iconic vocalizations can enable interlocutors to establish understanding through vocalizations in the absence of conventions, thereby demonstrating the iconic potential of speech. Contestants were able to create iconic vocalizations for a wide array of semantic domains, and these vocalizations were largely comprehensible to naïve listeners, as well as easier to learn as category labels. Along with other recent studies of iconicity in the production of vocalizations ^42,44^, including vocal imitation ^38^, our results complement the accumulating evidence of iconicity in the vocabularies ^24,32,46^, grammar ^47^, prosody ^37,39^, and in the acquisition^18,45^, of spoken languages. Taken together, these findings support the hypothesis that iconicity in human communication is not limited predominantly to gesturing and signed languages, but also plays an important role in our vocal communication, including speech ^17–20^. This newly emerging understanding of iconicity as a widespread property of spoken languages suggests iconicity may also have played an important role in their origin. An intriguing possibility is that many of the now arbitrary words in modern spoken languages may have originated from the innovation of iconic vocalizations.

## Methods

### Contest

#### Participants

We recruited contestants by advertising the contest on the Internet, including calls through www.replicatedtypo.com and through an announcement on the Protolang 4 conference website. We received 11 submissions: seven from individuals and four from teams. Nine of the submissions came from the United States, one from the United Kingdom, and one from Poland. There were two submissions by native German speakers, and one by native Polish speakers. Eight submissions came from researchers in relevant academic departments (e.g. Linguistics, Language Evolution, Psychology), and three from individuals not affiliated with academic institutions.

#### Stimuli

Our set of stimuli consisted of 30 basic meanings: 8 action verbs (*cook*, *cut*, *eat*, *gather, hide*, *hunt*, *pound*, *sleep*), 12 nouns referring to people (*child, man, woman*), animals (*snake, tiger, deer*), and inanimate things (*fire*, *fruit, knife*, *meat*, *rock*, *water*), and 10 properties, a heterogeneous category that included adjectives (*bad*, *big*, *dull*, *good*), quantifiers (*one, many*), and demonstratives (*that*, *this*). The 30 items included meanings in different categories that were semantically related and potentially confusable, e.g., *cook* and *fire* (*fire* is used to *cook*)*; cut, knife,* and *sharp* (*sharp knives* are used to *cut*); *rock*, *pound,* and *dull* (a *rock* is used to *pound*; a *rock* is *dull* compared to a *knife*); *small* and *child* (a *child* is *small*).

#### Submissions

Contest instructions and other details were made available at the original contest website: <u>http://sapir.psych.wisc.edu/vocal-iconicity-challenge/</u> and in the Supplementary Materials. Potential contestants were asked to submit a set of recorded vocalizations for each of the 30 meanings together with a brief statement explaining the rationale of each vocalization. Vocalizations were defined as sounds “produced by the vocal apparatus”. Contestants were permitted to produce imitative sounds, but not allowed to use recognizably onomatopoeic words or conventional emblems, e.g., “booo” (bad), “nyam nyam” (eat), or “roar” (the sound of a tiger). We verified that none of the submissions violated this rule in any clear way. The most marginal case was the use of sibilants for *snake*; however, in each instance, the sound was elaborated beyond any standard convention.

The instructions stipulated that each vocalization should be no longer than two seconds. Twenty-four (7.3%) of the submitted vocalizations exceeded this duration (16 from the last-placed contestant AB). The top contestant followed this restriction, and so it did not interfere with determining the winner of the contest. For completeness, we report results with all of the vocalizations included.

#### Determining the winner

The winner of the contest was determined by experiments that tested how well naïve listeners could guess the meaning of the vocalizations from each submission. We tested the vocalizations in two independent phases consisting of within-semantic category and between-semantic category testing (see below). As per the instructions, the top five submissions in the first phase of testing advanced to the second phase, and the submission with the highest overall guessing accuracy in the second phase was crowned the winner. In all of the analyses presented here, vocalizations from all 11 submissions were subjected to both testing conditions.

### Comprehensibility of vocalizations

#### Participants

We recruited 708 participants through Amazon Mechanical Turk to serve as listeners: 366 in within-category testing and 342 in between-category testing. We aimed to test each vocalization with 10 participants in each phase, but this number was sometimes inadvertently exceeded. Participants were restricted to be in the USA and were only permitted to participate once in the study.

#### Stimuli

The stimuli were the recorded vocalizations submitted by the contestants in the contest. Eleven contestants each produced vocalizations for 30 meanings, for a total of 330 vocalizations.

#### Design and procedure

Participants listened to a set of vocalizations from a single contest submission, and selected the meaning of each from a list of alternatives. They could listen to each vocalization as many times as they needed before making their choice. The vocalizations were presented in random order. To ensure that listeners properly attended to the task, we also included catch trials with the spoken phrase “cats and dogs”, along with a corresponding option. Four participants did not respond correctly to these catch trials and were excluded from further analysis.

The vocalizations of each submission were tested separately in two conditions (*within* and *between*). Figure 7 shows trial schematics from each testing condition. In the *within* condition vocalizations were presented with foils from within the same word class. Thus the meaning of a vocalization for an action was selected from 8 alternatives, a vocalization for a property from 10, and a vocalization for noun from 12. On *between* testing trials, the foils came from between all word classes. Meanings were placed into three lists of 10, with some thematically related meanings deliberately included to increase difficulty (e.g. *rock*, *pound*, *dull* and *knife, cut, sharp*; Figure 3 shows the meanings used in each list).

**Figure 7.**
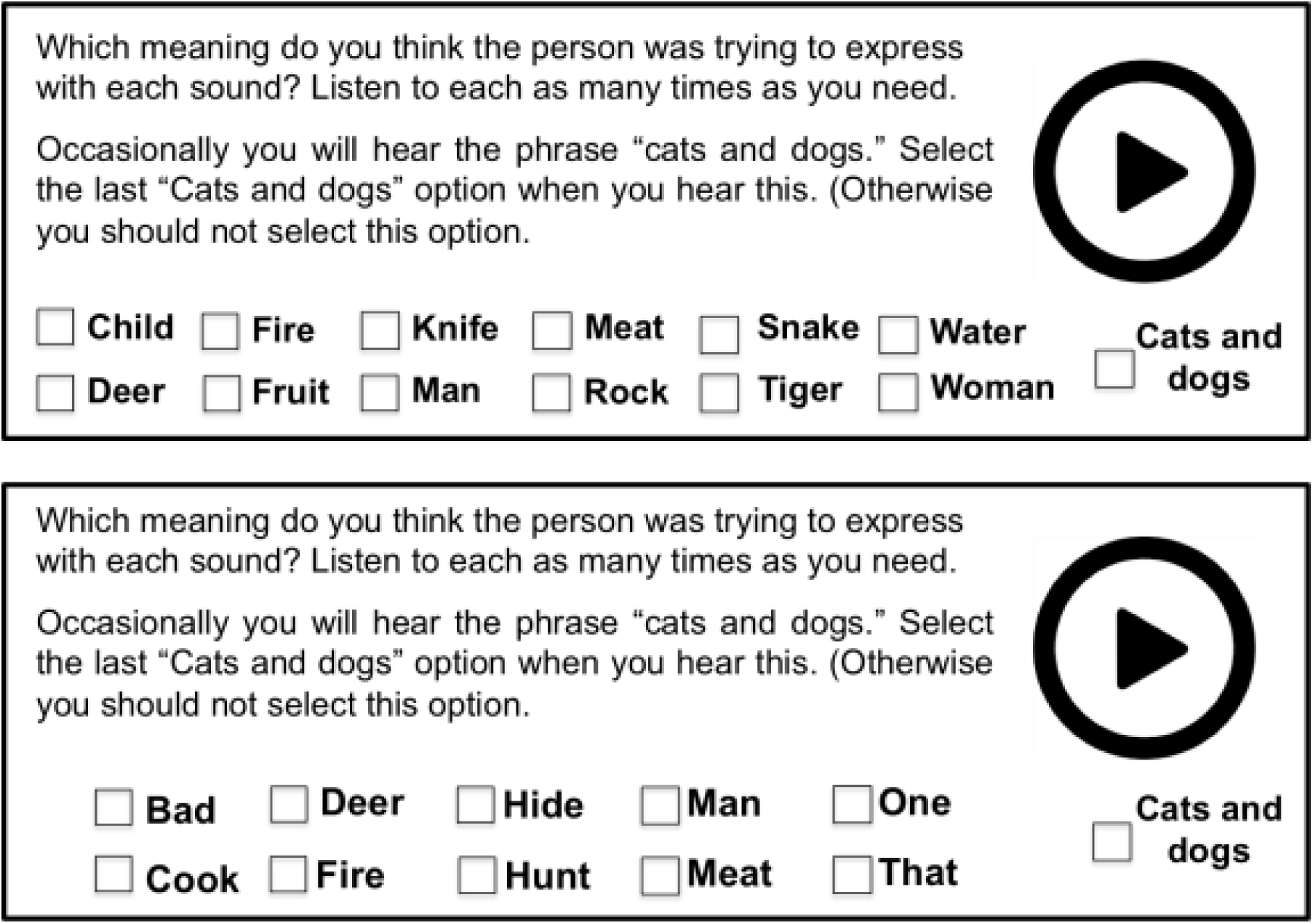
A schematic of the testing method. The top panel shows a within-category trial with nouns. The bottom panel shows a between-category trial.

### Measuring semantic similarity and confusability

To determine whether listeners were more likely to confuse similar meanings, we used Google’s *Word2Vec* semantic vectors to index the semantic similarity between pairs of words. Google’s *Word2Vec* is a model that produces vector-based representations of words such that words occurring in similar contexts are represented in similar ways^48,49^. This makes it possible to capture similarity relations between words that occur in similar contexts even if they never occurred in the same context ^50,51^. Specifically, *Word2Vec* is a three-layer neural network that is trained by being presented one word at a time; its task is to predict the words that surround the input word. Although differing in details, the logic of this predictive model is broadly similar to that of Simple Recurrent Networks^52^, which, when trained on a small word corpus are able to extract lexical classes (nouns, verbs) and broad semantic fields (animals, edible items, etc.)^53^. Models like *Word2Vec* are able to capture a considerable amount of variance of human word-to-word similarity ratings^54–56^, in part due to being trained on vastly larger datasets such as Google News.

We used the correlation between the *Word2Vec* vectors of pairs of words to indicate the degree to which those words occur in similar contexts, ranging from 0 (most different) to 1 (most similar). For each vocalization in each of the two testing conditions, we computed the proportion of trials in which the intended meaning was matched with each possible alternative. In this way, each vocalization, with its intended meaning, generated a series of data points – including one point for each alternative meaning in within-category testing, and one point for each alternative in between-category testing – that index how frequently that particular vocalization was matched with each possible alternative (including the correct meaning). For example, in between-category testing, the vocalization of one submission for “sharp” was guessed to be “cut” in 20% of trials, ‘good’ in 70% of trials, ‘snake’ in 10% of trials, as the correct response ‘sharp’ in 0% of trials, and likewise, the remaining six alternatives were selected in 0% of trials. The *Word2Vec* similarity index for these combinations was 0.24 between ‘sharp’ and ‘cut’, 0.30 between ‘sharp’ and ‘good’, 0.11 between sharp and ‘snake’, etc. (and 1 with itself).

### Ratings of form-meaning resemblance

To assess the degree to which listeners perceived a resemblance between the forms of the vocalizations and their intended meanings, we asked naïve listeners to directly rate the degree to which each one “sounds liked what it means”.

#### Participants

We recruited 229 new participants through Amazon Mechanical Turk. The target was to acquire 10 ratings for each vocalization (i.e. 10 raters for each of the 22 lists), and this number was exceeded in a few cases. Participants were restricted to be in the USA and were only permitted to participate once in the study.

#### Stimuli

The stimuli were the 330 recorded vocalizations submitted by the 11 contestants in the contest.

#### Design and procedure

The vocalizations were separated into 22 pseudo-randomized lists of 15 different meanings. Each list contained vocalizations from each of the 11 contestants, with 4 contestants repeated twice.

Using a procedure similar to ^45^, we collected ratings of the iconicity of the vocalizations. Participants listened to the vocalizations in a random order, with the meaning of each one displayed. For each vocalization, they were asked to indicate “how much it sounds like what it means” on a scale of −5 (very opposite) to 0 (arbitrary) to 5 (very iconic).

**Figure 8.**
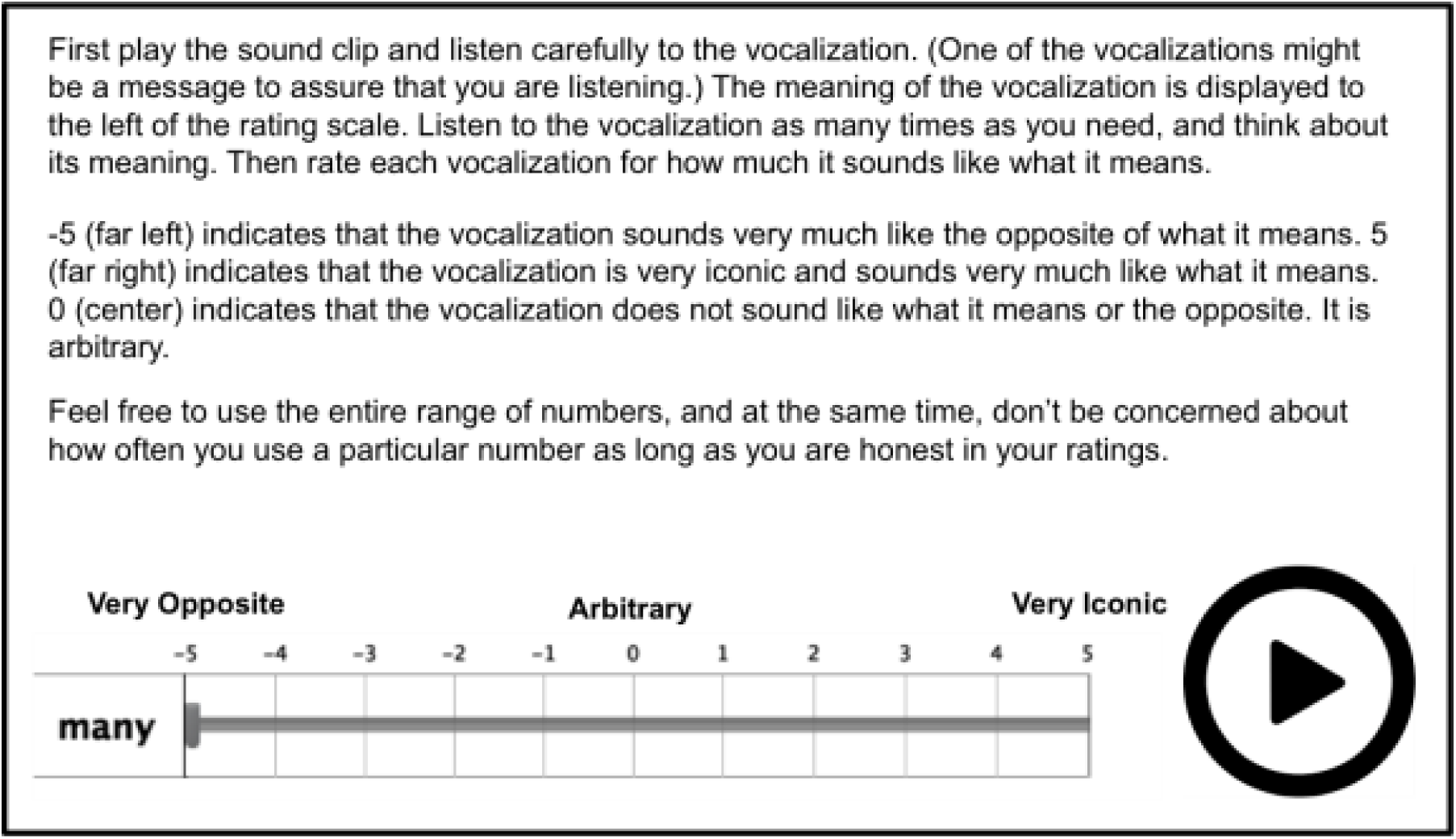
Schematic of the interface used for iconicity ratings.

### Agreement between producers and listeners

To measure the degree to which contestants produced similar vocalizations for each meaning, we devised a similarity metric based on the acoustic properties of the vocalizations. We used Praat phonetic analysis software (Boersma, 2001) to measure the duration, pitch, intensity, and harmonics to noise ratio of the vocalizations. The onset and offset of each vocalization was marked by hand, and a script was used to automatically measure the duration, intensity, and harmonics to noise ratio. Pitch was measured by hand using the pitch tracker to avoid spurious measurements that can be common, such as doubling or halving of pitch, especially in weakly voiced onsets and offsets of the vocalizations. The formula for the similarity metric is shown below. For each meaning (M), the standard deviation of each property (DUR = duration, PIT = pitch, INT = intensity, HNR = harmonics to noise ratio) was divided by the mean value of that property over all meanings. These four values were then added together, providing a metric of overall similarity of the vocalizations for each meaning.

~~~
SIM.DUR_M_ = SD.DUR_M_ / MEAN.DUR_Total_
~~~

~~~
SIM.PIT_M_ = SD.PIT_M_ / MEAN.PIT_Total_
~~~

~~~
SIM.INT_M_ = SD.INT_M_ / MEAN.INT_Total_
~~~

~~~
SIM.HNR_M_ = SD.HNR_M_ / MEAN.HNR_Total_SIM_M_
~~~

~~~
= SIM.DUR_M_ + SIM.PIT_M_ + SIM.INT_M_ + SIM.HNR_M_
~~~

### Learnability of vocalizations as category labels

#### Participants

We recruited 87 undergraduate students from University of Wisconsin-Madison to participate in exchange for course credit.

#### Stimuli

The stimuli consisted of a subset of the recorded vocalizations from the contest.

Owing to the difficulty of depicting actions and properties with static images, we restricted our materials to the 12 nouns. For each of the 12 nouns, we selected 10 different pictures that clearly depicted its referent (e.g. 10 pictures of a fire for *fire*, 10 pictures of fruit for *fruit*).

For each meaning, three vocalizations were selected on the basis of the iconicity ratings. These included the vocalization with the lowest mean iconicity rating, the one with the median rating, and the one with the highest rating. These comprised the low, medium, and high iconicity conditions, respectively.

#### Design and procedure

Participants were randomly assigned to one of the three iconicity conditions and, within each condition, one of two feedback conditions as described below. On each trial, participants heard a vocalization and attempted to select its meaning from 12 pictures (arranged in a 3×4 grid) representing each of the different meanings. Participants completed 10 blocks with each block including vocalizations from each of the 12 categories, in a random order. The position of the images and the specific exemplar used for the meaning were randomized with the stimulation that all category exemplars of a target category were used as targets at some point for each participant. Participants selected the image depicting the category of the vocalization made their selection by clicking on one of the pictures with the mouse. They then received feedback on their selection according to one of two randomly assigned conditions. In the *full feedback* condition, the correct response was explicitly indicated by highlighting the correct image. In the *accuracy only* condition, the participant was informed only whether their response was correct or incorrect. Because each image served as a target exactly once throughout the experiment, participants could not respond correctly simply by associating a sound with a specific image.

### Statistical analyses

Statistical analyses with mixed effects models were conducted using the lme4 package version 1.1-10 ^57^ in R version 3.2.3 ^58^. Significance tests of continuous outcomes were calculated using χ2-tests that compared the model likelihoods with and without the factor of interest. Significance tests of binomial outcomes used the z-values associated with the logistic mixed-effect models. To compare outcomes to chance levels, the logit function – the logarithm of the odds – was applied to the chance level, and this was included in the model as an offset to the intercept. Thus, an intercept greater than zero indicated that the outcome exceeded the level of chance.

### Ethics

All experiments were approved by the Institutional Review Board of the University of Wisconsin-Madison, and were conducted according to the relevant guidelines and regulations. All participants provided informed consent as required according to the approved experimental protocols.

### Data availability

Data and analysis scripts, including vocalizations from the contest, are available through the Open Science Framework at osf.io/x9w2z.

## Authors’ Contributions

M. Perlman and G. Lupyan devised the experiments, analyzed the data, and wrote the manuscript.

## Competing Interests

We have no competing interests.

## Funding

This work was funded in part by an NSF INSPIRE Award 1344279 to GL.

